# An *ex vivo* model of interactions between extracellular vesicles and peripheral mononuclear blood cells in whole blood

**DOI:** 10.1101/2023.05.11.540421

**Authors:** Blanca V. Rodriguez, Yi Wen, Erin N. Shirk, Samuel Vazquez, Olesia Gololobova, Amanda Maxwell, Jessica Plunkard, Natalie Castell, Bess Carlson, Suzanne E. Queen, Jessica M. Izzi, Tom A.P. Driedonks, Kenneth W. Witwer

**Affiliations:** Department of Molecular and Comparative Pathobiology, Johns Hopkins University School of Medicine, Baltimore, MD, USA; University Medical Center, Utrecht University, Utrecht, The Netherlands; Department of Neurology, Johns Hopkins University School of Medicine, Baltimore, MD, USA; Richman Family Precision Medicine Center of Excellence in Alzheimer’s Disease, Johns Hopkins University School of Medicine, Baltimore, MD, USA

## Abstract

Extracellular vesicles (EVs) can be loaded with therapeutic cargo and engineered for retention by specific body sites; therefore, they have great potential for targeted delivery of biomolecules to treat diseases. However, the pharmacokinetics and biodistribution of EVs in large animals remain relatively unknown, especially in primates. We recently reported that when cell culture-derived EVs are administered intravenously to *Macaca nemestrina* (pig-tailed macaques), they differentially associate with specific subsets of peripheral blood mononuclear cells (PBMCs). More than 60% of CD20^+^ B cells were observed to associate with EVs for up to 1 hr post-intravenous administration. To investigate these associations further, we developed an *ex vivo* model of whole blood collected from healthy pig-tailed macaques. Using this *ex vivo* system, we found that labeled EVs preferentially associate with B cells in whole blood at levels similar to those detected *in vivo*. This study demonstrates that *ex vivo* blood can be used to study EV-blood cell interactions.

## Introduction

Extracellular vesicles (EVs) are membranous nanosized particles released constitutively into the extracellular space by virtually all cells. EVs carry proteins, nucleic acids, metabolites, and lipids reflective of the contents of the producing cells and modulate physiological and pathological processes of recipient cells ^1–5^. Although EVs are thought to have promise in therapeutic applications ^1,6–8^, a deeper understanding is needed of the pharmacokinetics and biodistribution of EVs in animal models beyond rodents ^9–18^. Rodent models of human disease are valuable but sometimes respond to experimental interventions in markedly different ways from humans ^19^.

Recent studies have begun to uncover the pharmacokinetics and biodistribution of EVs in larger animals, including in non-human primates ^20–23^. We recently reported that when EVs isolated from cell culture are administered intravenously to *Macaca nemestrina* (pig-tailed macaques), they differentially associate with specific subsets of peripheral blood mononuclear cells (PBMCs) ^23^. Intriguingly, flow cytometry identified CD20^+^ B cells as the predominant immune cell population capable of interacting with EVs, and this finding was consistent for EVs that were labeled genetically (with GFP) and post-production with a self-quenching lipid dye.

In this study, we sought to establish an *ex vivo* model to further interrogate EV-blood cell associations. We reasoned that an *ex vivo* model would have several advantages to *in vivo* studies, including precise control of experimental conditions; minimal invasiveness, i.e., only one donor intervention (single blood draw) rather than administration and repeated sampling; and applicability to donors of multiple species, including humans. Flow cytometry and nanoluciferase assays tracked the distribution of labeled EVs in the plasma, PBMCs, and red blood cell compartments of *ex vivo* blood. Here, we show that EVs derived from both Expi293F and U-87 MG cells disseminate to all blood compartments but appear to interact preferentially with CD20^+^ B cells. This study demonstrates that *ex vivo* blood can be used to study EV-blood cell interactions recapitulating *in vivo* results.

## Materials and Methods

### Cell culture and EV separation and labeling

Expi293F cells were cultured and transiently transfected with the pLenti-palmGRET reporter plasmid encoding the dual reporter palmitoylated EGFP-nanoluciferase protein (PalmGRET) (Addgene plasmid #158221 ^11^) for EV production as described previously ^23^. EVs were separated from the conditioned medium three days post-transfection as follows: PalmGRET-transfected Expi293F cells were centrifuged at 1,000 × g for 20 min at 4 °C, the collected supernatant was then centrifuged at 2,000 × g for 20 min and filtered through 0.22 μm bottle-top filters (Corning, NY). EVs were then aditionally separated and concentrated 10X by tangential flow filtration (TFF, Sartorius Vivaflow 100 kDa MWCO). Some TFF-concentrated EVs were fluorescently labeled with 200 nM MemGlow 700 nm dye (Cytoskeleton, catalog #MG05-10) for 30 min at room temperature (RT) in the dark. EVs were further concentrated and unincorporated MemGlow 700 dye was removed by centrifugation with Amicon 15 Ultra RC 10k Da MWCO filters (EDM Millipore). PalmGRET EVs not labeled with MemGlow 700 were also further concentrated after TFF on Amicon 15 Ultra RC 10 kDa MWCO filters. Filtered EVs were further purified using qEV10 70 nm Legacy size-exclusion chromatography (SEC) columns (Izon). EV-enriched fractions (1-4) were pooled together, concentrated on Amicon 15 Ultra RC 10 kDa MWCO filters, and aliquoted/stored in Dulbecco’s phosphate buffered saline (DPBS) at -80 °C in protein LoBind tubes (Eppendorf).

U-87 MG cells (ATCC-HTB-14) were maintained in Dulbecco’s modified Eagle’s medium (DMEM; Gibco) supplemented with 10% fetal bovine serum (FBS, heat-inactivated), 2 mM L-glutamine (Gibco, catalog #25030081), 10 mM HEPES (Gibco, cat #15630106), 100 U/mL penicillin –streptomycin (Gibco, cat #15140122). U-87 MG cells were grown in a humidified 5% CO_2_ incubator set to 37 °C. For EV isolation, cells were incubated in EV-depleted media (EV-depleted FBS, Gibco, catalog #A2720801). Cell conditioned media (CCM) was collected after 48 hours of harvesting. Cells were removed by centrifugation at 300 × g for 5 min at 4 °C, supernatant were further centrifuged at 2,000 × g for 10 min at 4 °C. EVs were separated by ultracentrifugation (UC) or SEC. UC-EVs were obtained by centrifuging 50 mL of conditioned media at 100,000 × g for 70 min at 4 °C (AH-629/36 rotor, Beckman Ultra-Clear Tubes with 36 ml capacity). Pellet was re-suspended in 250 µL of DPBS. For SEC, CCM was concentrated with an Amicon 15 Ultra RC 10 kDa MWCO filters from 50 ml to 0.5 ml before application onto qEV Original 70 nm Legacy SEC column. EV-enriched fractions (1-4) were pooled and concentrated with Amicon 4 Ultra RC 10 kDa MWCO filters to 250 μl.

For MemGlow labeling, U-87 EVs were adjusted to 1E+10 particles/mL in PBS and stained with 200 nM MemGlow 700 nm at RT for 30 minutes, protected from light. Excess dye was removed by ultrafiltration as above, and labeled EVs were stored overnight at 4 °C prior to whole blood experiments.

### EV quantification by nano flow cytometry

Particle concentration, size, and %GFP positivity of EV preparations were measured with a NFCM Flow NanoAnalyzer (NanoFCM Co., Ltd) following the manufacturer’s instructions and as previously reported ^24^. Briefly, lasers were aligned and calibrated separately for particle concentration using fluorescent 250 nm silica nanoparticles at a concentration of 2.19E+10 (NanoFCM, catalog #QS2503) and for size using a premixed silica nanosphere cocktail containing monodisperse nanoparticle populations of 68 nm, 91 nm, 113 nm, and 155 nm in diameter (NanoFCM, catalog #516M-Exo). DPBS was used as the blank for background correction. For quantification of EVs using NFCM, MemGlow labeled and unlabeled PalmGRET-EV preparations were diluted 50-fold or 10,000-fold in DPBS, respectively, and U-87-derived EVs were diluted 100-fold in DPBS. Particle signal acquisition was performed for 1 min at constant pressure of 1 kPa at an event rate between 1,500 and 10,000 events/min. The side-scattering signal and fluorescent signal of each sample were calculated using the NanoFCM Professional Suite V2.0 software.

### Western blot characterization of U-87 MG EVs

18 µL U-87 MG EVs that had been separated by UC or SEC were lysed in 1x radioimmunoprecipitation assay buffer (RIPA, Cell Signaling Technology, catalog #9806) for 30 minutes at room temperature. Lysates were heated at 95 °C for 5 min together with Laemmli sample buffer (Bio-Rad, catalog #1610747). Lysates were resolved using a 4% to 15% Criterion TGX Stain-Free Precast gel (Bio-Rad, catalog #5678084), with Spectra Multicolor Broad Range protein ladder (Thermo Scientific, catalog #26634), then transferred onto a PVDF membrane (Invitrogen, catalog #IB24001) using iBlot 2 semi-dry transfer system (Invitrogen) for 1 minute at 20 V, 4 minutes at 23 V, and 2 minutes at 25 V. Blots were probed using primary antibodies in PBST (1X DPBS with 0.05% Tween-20 (BioXtra, catalog #P7949) with 5% Blotting Grade Blocker (Bio-Rad, catalog #1706404) for approximately 16 hours (overnight) at 4 °C. The following primary antibodies were used: mouse-anti-CD63 (BD Biosciences, catalog #556019, dilution (dil) 1:1000), mouse-anti-CD9 (Biolegend, catalog #312102, dil 1:1,000), rabbit-anti-ALIX (abcam, catalog #ab186429, dil 1:1000), rabbit-anti-Calnexin (abcam, catalog #ab22595, dil 1:2,000), mouse-anti-Bovine Albumin (Invitrogen, catalog #MA515238, dil 1:500). After washing in PBST-milk four times, blots were incubated with corresponding secondary antibodies: mouse-IgGk BP-HRP (Santa Cruz Biotechnology, catalog #sc-516102, dil 1:10,000) or mouse anti-rabbit IgG-HRP (Santa Cruz Biotechnology, catalog #sc-2357, dil 1:10,000) for 1 hour in PBST-milk. After two washes in PBST-milk and two washes in PBST, blots were incubated for 30 seconds in SuperSignal West Pico PLUS Chemiluminescent Substrate (Thermo Scientific, catalog #34580) and visualized with an iBright 1500FL Imager (Thermo Fisher, Waltham, MA).

### EV spike-in experiments with *ex vivo* blood

Up to 20 mL of whole blood per donor was drawn by venipuncture into a sterile 50-mL Luer-lock syringe tube (BD catalog #309653) containing acid citrate dextrose (ACD) at an ACD-to-whole blood ratio of 1:5. Blood was processed within 1 hour of collection. Under sterile conditions in a cell culture hood, whole blood was transferred to a separate 50 mL conical tube and further aliquoted into sterile 2-mL or 5-mL Eppendorf tubes. For time-course experiments, 1 mL of whole blood was incubated with 8E+08 MemGlow 700-labeled PalmGRET-EVs or an equal volume of DPBS for up to 24 hours in an incubated rotator at 37 °C, with the “Mix” mode selected at a speed of 8 rpm (Benchmark Scientific Roto-Therm Plus Incubated Rotator, item #H2024). Dose-response assays were conducted by incubating 4 mL of whole blood with 1.4E+08, 8E+08, 4E+09, or 1.8E+10 PalmGRET-EVs for 30 minutes. Dosages were established to (1) roughly match those used in our previous *in vivo* studies; (2) allow for detection of signal above background after processing; and (3) avoid the input exceeding 10% of total volume. One tube containing whole blood incubated with DPBS of volume equal to 2E+10 EVs served as the treatment control group. Tubes were removed from the rotating incubator at each timepoint, and 100 µL of blood was set aside for flow cytometry analysis. The remaining blood was processed as described below for nanoluciferase activity assays.

### Flow cytometry

To quantify the association between PBMCs and PalmGRET EVs in whole blood, PBMCs were immunolabeled directly in whole blood with fluorescent antibodies. Following incubation of whole blood with MemGlow-PalmGRET-EVs, PalmGRET-EVs, U-87-EVs, or DPBS, 100 µL of whole blood was added to an antibody cocktail containing the following antibodies: mouse-anti-CD3-V500 (BD, catalog #560770, dil 1:30), mouse-anti-CD4-PerCP/Cy5.5 (BD Biosciences, catalog #552838, dil 1:7.5), mouse-anti-CD8-BV570 (BioLegend, catalog #301038, dil 1:60), mouse-anti-CD20-e450 (Thermo Fisher, catalog #48-0209-42, dil 1:60), mouse-anti-CD159a-PE (Beckman Coulter, catalog #IM3291U, dil 1:30), and mouse-anti-CD14-BV650 (BioLegend, catalog #563419, dil 1:30). The mixture was briefly vortexed and incubated at RT for 20 min. Red blood cells were then lysed for 10 min at RT with 2 mL RBC lysis buffer composed of 0.83% NH_4_Cl, 0.1% KHCO_3_, and 0.03% ethylenediaminetetraacetic acid, EDTA, and pre-warmed to 37°C. Lysed RBCs were removed by centrifugation at 400 × g for 5 minutes and PBMCs were washed once with PBS. Pelleted PBMCs were resuspended in 500 µL PBS, and PBMC-associated EGFP and MemGlow 700 signal were measured directly on a BD LSR Fortessa flow cytometer. Fluorescence minus one (FMO) controls for CD159a and CD4 were used for accurate gating of GFP, PE and PerCP/Cy5.5 fluorescence.

### Fractionation of EV-treated whole blood

Different blood compartments (plasma, PBMCs, and RBCs) were separated by density gradient centrifugation using SepMate-15 tubes (STEMCELL Technologies, catalog #85415) to reduce blood processing time and user variability, as described previously ^25^. Each whole blood sample was diluted 1:1 in PBS-EDTA (1 mM) + 2% FBS buffer (PBS-EDTA-FBS buffer) in a 15-mL conical tube and mixed well. Diluted blood was overlaid carefully on SepMate-15 tubes each containing up to 4 mL Lymphoprep (STEMCELL Technologies, catalog #07851). The layered blood was centrifuged at 1,200 × g for 10 minutes at RT, with the centrifuge brake on maximum settings. After centrifugation, 5 mL of plasma from the top layer of each sample was transferred to sterile 15-mL conical tubes and stored at -80 °C for later downstream analysis. The remaining upper plasma layer was discarded without disturbing the Lymphoprep-plasma interface.

PBMCs were collected from the Lymphoprep-plasma interface layer into 15-mL conical tubes, washed with up to 10 mL PBS-EDTA-FBS buffer, and centrifuged at 400 × g for 8 minutes at RT. PBMC pellets were resuspended in 1 mL RBC lysis buffer and incubated for 5 min at 37°C to deplete contaminating RBCs from PBMCs. PBMCs were then washed once more after RBC lysis by adding up to 10 mL PBS-EDTA-FBS followed by centrifugation for 8 min at 400 × g at RT. PBMC pellets were resuspended in 0.5 mL of cell freezing media (90% FBS/ 10% DMSO), transferred to cryovials, and placed in a Mr. Frosty freezing container (Thermo Scientific, catalog #5100-0001) with isopropanol at -80 °C. After at least 12 h in the -80°C freezer, PBMC-containing cryovials were transferred to a liquid nitrogen tank for long-term storage. For analysis, PBMCs were rapidly thawed in a 37 °C water bath with constant swirling, transferred to a 15-mL conical tube containing 8 mL of warm RPMI 1640 + 10% FBS media, recovered by centrifugation at 400 × g for 8 minutes, resuspended in 200 µL lysis buffer (PBS + 1% Triton X-100 + 1 Complete Mini protease inhibitor tablet), and kept on ice for 15 min. Lysates were centrifugated at 16,000 × g for 15 min at 4 °C to spin out cell debris, and the supernatant was collected for protein and nanoluciferase assays as described below.

RBC lysates were prepared by transferring 1 mL of RBC sample from the bottom of each SepMate-15 tube to a 15-mL conical tube. RBCs were lysed with 1 mL of RBC lysis buffer for 5 minutes at 37°C, and RBC lysates were stored directly at -80 °C.

### Nanoluciferase assays

Nanoluciferase assays were used to quantify the presence of PalmGRET-EVs. 50 µl undiluted plasma, RBC lysate, and PBMC lysate samples were loaded in duplicate into a 96-well white flat bottom polystyrene plate (Corning, catalog #3922). Nanoluciferase activity was measured using the Nano-Glo Luciferase Assay System (Promega, catalog #N1110) per the manufacturer’s instructions. Briefly, Nano-Glo luciferase assay reagent was prepared just before use by combining one volume Nano-Glo luciferase assay substrate (furimazine) with 50 volumes Nano-Glo luciferase assay buffer. 50 μL Nano-Glo luciferase assay reagent was added to wells containing samples, and nanoluciferase activity was measured immediately on a BioTek Synergy 2 microplate reader in luminescence mode, integration time 20 ms. Results were normalized by total protein concentrations as determined by Pierce BCA Protein Assay Kit (ThermoFisher Scientific, catalog #23225). PBMC lysates were not diluted for this assay; plasma and RBC lysates were diluted 12.5-fold in PBS.

### Statistical analysis

Experimental replicates are defined in the figure legends for each experiment. Flow cytometry data were analyzed with FlowJo software (v10.8). Statistical analyses of flow cytometry data were performed with GraphPad Prism 9.0 (GraphPad Software Inc.) using multiple comparison analysis testing in two-way analysis of variance (ANOVA) with Tukey’s post-hoc test, or two-tailed paired t-tests, with *P* < 0.05 considered statistically significant.

## Results

We used *ex vivo* whole blood to reproduce and extend our unexpected finding that intravenously administered EVs associate at high levels with B cells in primates ^23^. Our experimental design, using whole blood from donor subjects and maintained at 37 °C, matched the conditions of the *in vivo* study as much as possible (Figure 1). Expi293F-derived EVs that were introduced into blood were labeled genetically with the dual reporter palmitoylated EGFP-nanoluciferase protein (PalmGRET) ^11^ and/or post-production with the near-infrared, self-quenching membrane dye MemGlow 700 (Figure S1A).

**Figure 1.**
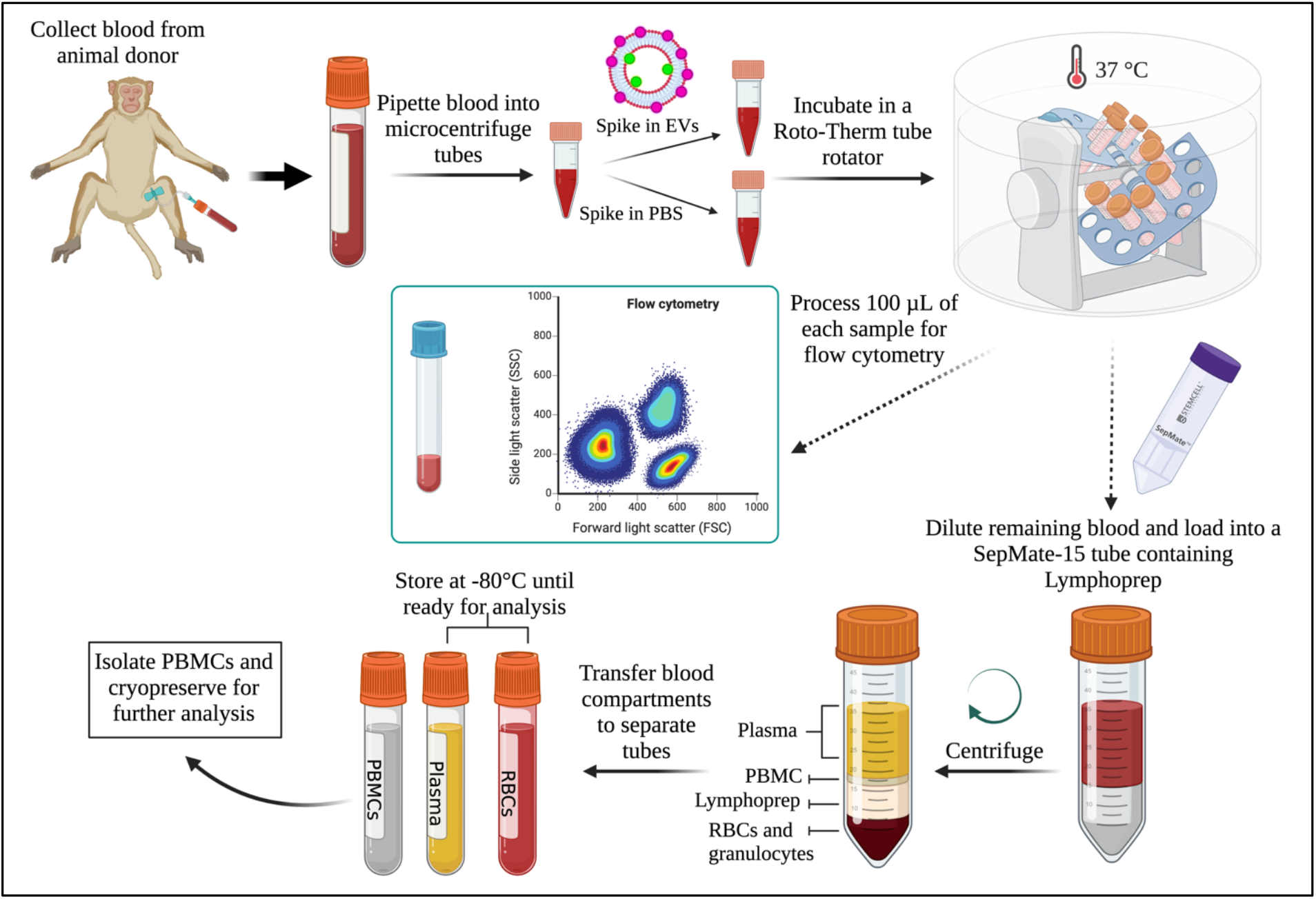
Schematic overview of the study design. Whole blood from pig-tailed macaques was drawn directly into a collection tube containing ACD. Whole blood was transferred to microcentrifuge tubes for EV spike-in time course and dose-response assays. Tubes were removed from the rotating incubator at predefined time points, and 100 µL of blood was set aside for PBMC staining and flow cytometry analysis. The remaining blood was divided into plasma, red blood cell lysates, and PBMCs using density gradient centrifugation. Once separated, each whole blood component was stored as indicated until analysis by nanoluciferase assays. Created with BioRender.com.

### Characterization of MemGlow-PalmGRET EVs

MemGlow-PalmGRET EVs were characterized for particle concentration, size, and % GFP positivity (Figure S1B, C). By nano-flow cytometry analysis, EV preparations contained >50% GFP+ particles, consistent with our previous results ^23^. 22 two-fold serial dilutions of EVs, starting from a concentration of 2.0E+10 particles/mL, were measured by nanoluciferase assay to assess linear range (Figure S2A). Since correlation between particle input and relative light units (RLU) weakened below 6.5 RLU or 1.64E+06 particles/mL, we considered signal below 6.5 RLU to be background signal throughout this study (Figure S2B). Please note that the particle counts here, as established by NFCM in our laboratory, may not be comparable with particle counts obtained in other laboratories or with other methods ^26^.

### Stability of EV-associated nanoluciferase in compartments of whole blood

We performed a 24-h time course to test the stability and distribution of nanoluciferase signal from EVs spiked into whole blood *ex vivo*. After fractionating blood treated with EVs or into plasma, RBC, and PBMC compartments, nanoluciferase activity remained detectable in each compartment for 24 h after spike-in (Figure 2). In contrast with our *in vivo* findings ^23^, nanoluciferase activity remained relatively stable over 24 hours in plasma and PBMC compartments of *ex vivo* blood (Figure 2A, C, D). Signal was comparatively low in the RBC compartment and declined steadily (Figure 2B). There was consistently more signal in plasma than PBMCs at each time point (Figure 2D), although not always statistically significant due to variability.

**Figure 2.**
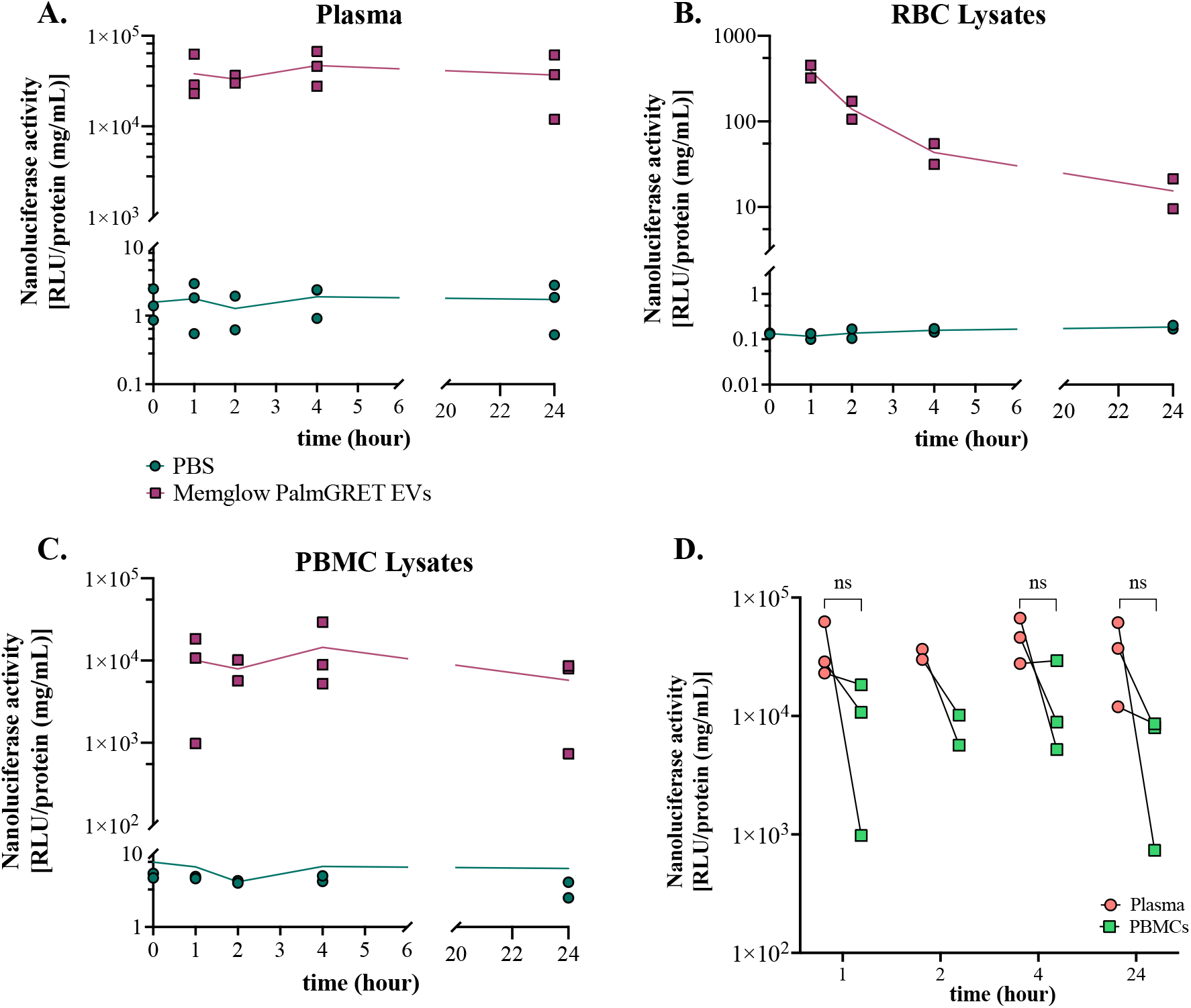
Distribution of EVs in macaque whole blood compartments at different time points *ex vivo*. PalmGRET EVs were detected by nanoluciferase assay in (A) plasma, (B) RBCs, and (C) PBMCs after addition of 8E+08 EVs or PBS. Nanoluciferase activity was normalized by protein concentration (BCA). Curves: mean of replicates (n=3 or n=2 donors as indicated for 2 h). (D) Comparison of nanoluciferase activity in plasma and PBMC compartments, normalized by total protein concentration. Each line represents a separate donor. Data points: results of two or three independent experiments, performed with two or three independent donors. Two-tailed paired t-tests were used to compare plasma and PBMC samples in D (p < 0.05 was considered significant).

### Time course analysis of EV interactions with PBMC subtypes

To determine which populations of PBMCs interact with EVs in whole blood *ex vivo*, subsets of PBMCs in whole blood were labeled with a fluorescent antibody panel at 5 min, 1 h, 2 h, 4 h, or 24 h after spike-in of 8E+08 EVs or PBS. See Figures S3 and S4 for cell and GFP+ and/or MemGlow+ gating. In line with our *in vivo* observations ^23^, EV signal was strongly associated with B cells across all time points (Figures 3A, 5A, and S4A). Around 80% of CD20^+^ B cells were positive for both genetic label and dye label at early time points, with signal declining slightly by 24 h (Figure 3A). Lower levels of signal were found in association with the other PBMC subtypes, and predominantly at early time points (Figure 3B-F; see also Figure 5B-F). Figure S4 shows the distribution of GFP and MemGlow in the form of overlaid dot plots and histograms. Of potential interest, although MemGlow and GFP signals were concordant in CD20^+^ B cells, this was not the case for every PBMC subtype. For example, MemGlow signal in CD3^+^ T cells and CD3^+^CD4^+^ T cells at 1 h was lower than GFP+ signal (Figure 3B, C), while the opposite was observed for CD159a NK cells and monocytes (Figure 3E, D).

**Figure 3.**
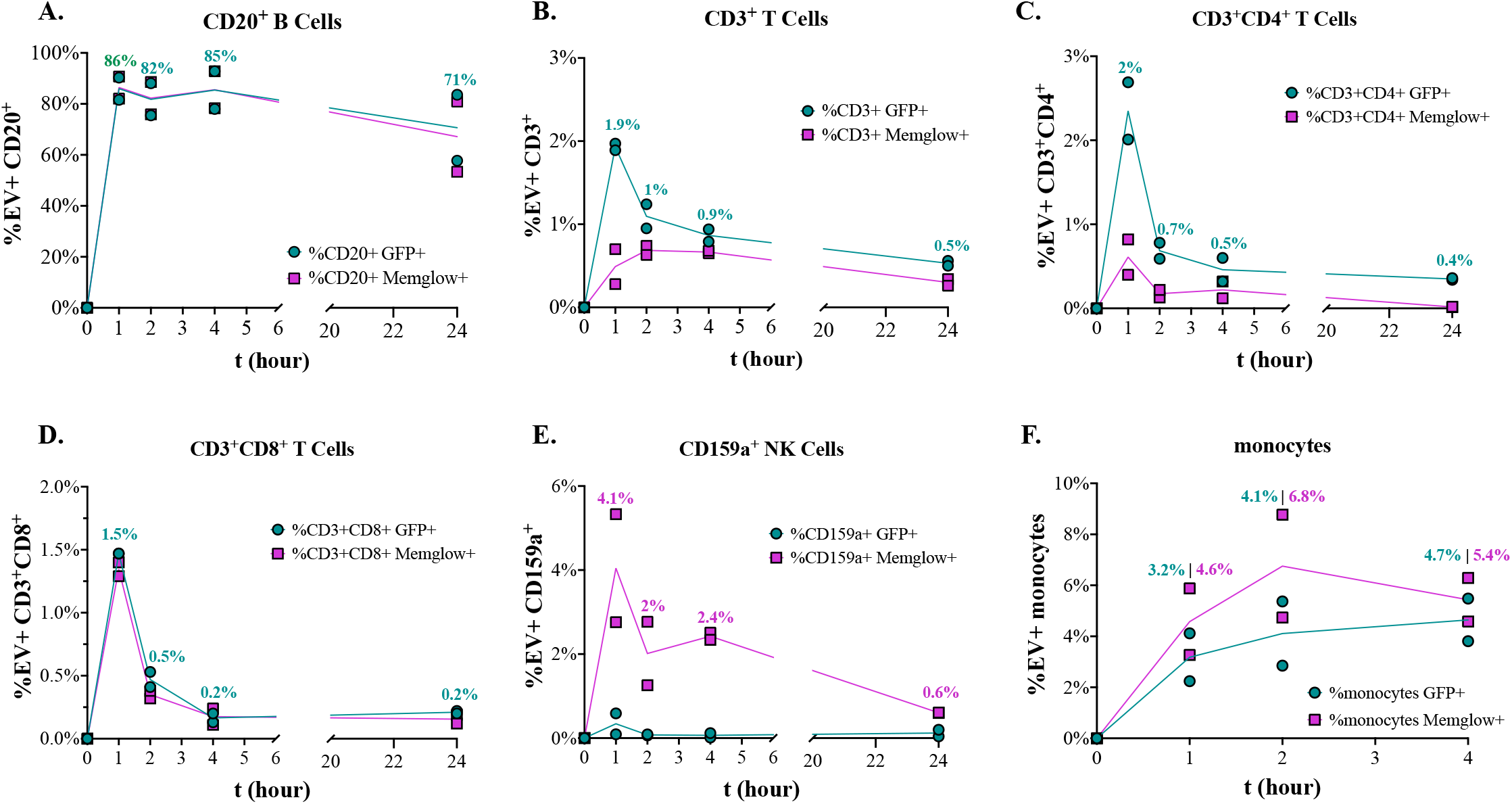
PalmGRET EVs preferentially interact with CD20^+^ B cells over a 24 h time course. GFP and MemGlow signals were detected by flow cytometry for (A) CD20^+^ B cells (B) CD3^+^ T cells, (C) CD3^+^CD4^+^ T cells, (D) CD3^+^CD8^+^ T cells, (E) CD159a^+^ NK cells, and (F) monocytes at time points over up to 24 h after spiking 8E+08 MemGlow-PalmGRET EVs into whole blood. Data are expressed as % GFP+ or %MemGlow+ for each PBMC subtype. Data are from experiments with blood from n = 2 donors.

Prior to spike-in (t = 0), CD3^+^ T cells, CD20^+^ B cells, and monocytes accounted for 62%, 11%, and 8% of PBMCs, respectively, and recorded percentages remained roughly the same across time points after spike-in: 59-63% CD3^+^ T cells, 7-10% CD20^+^ B cells, and 1-8% monocytes (Figure 4A). These PBMC frequencies are normal for non-human primate blood ^27^ and did not change markedly after EV spike in (Figure 4A). From 1 h to 24 h post-spike, 8% to 11% of total PBMCs were GFP+. At all time points, CD20^+^ B cells were the majority of GFP+ PBMCs, followed by CD3^+^ T cells (Figure 4B).

**Figure 4.**
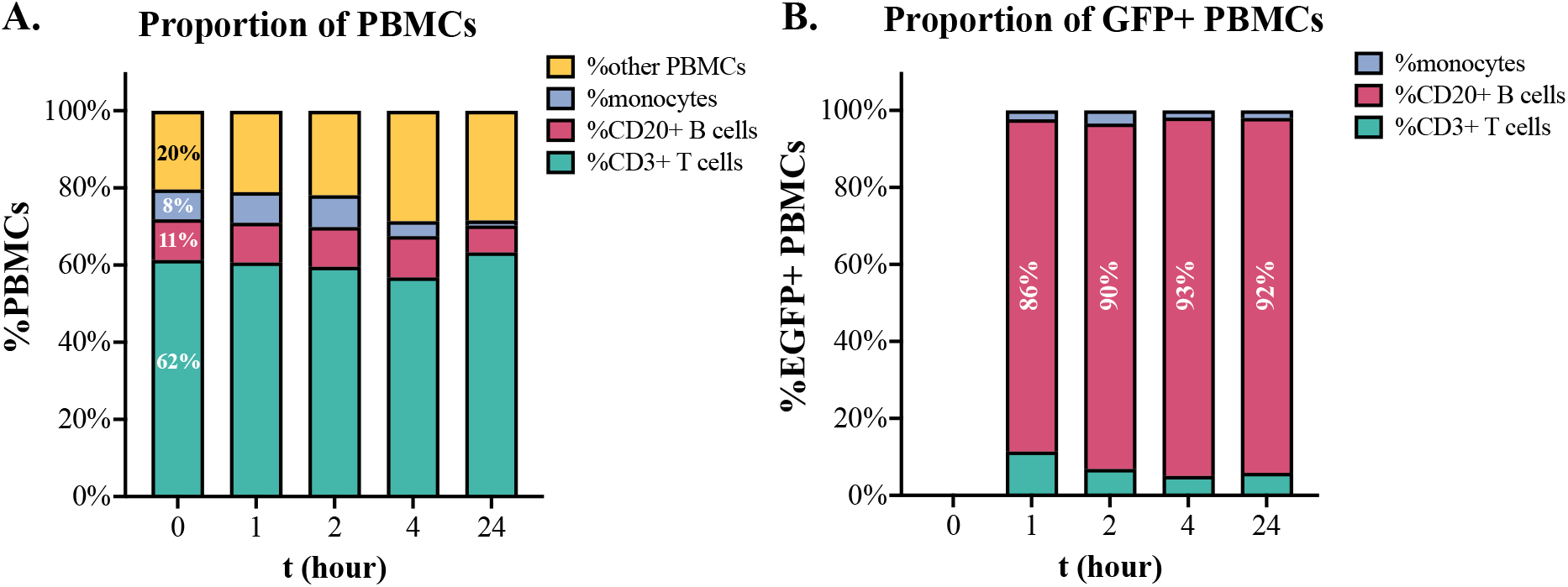
Specific subsets of PBMCs and their contribution to EGFP+ PBMCs. (A) Comparison of PBMC subtype proportions and (B) proportion of GFP+ PBMC subsets (CD3^+^ T cells, CD20^+^ B cells, monocytes). Graphs display mean frequency as a percentage of total PBMCs gated by PBMC subsets or GFP fluorescence. Data are from experiments with blood from n = 2 donors. B cells are a relatively small population of total PBMCs but are the majority of detectable GFP+ PBMCs.

### Effect of EV source and separation strategy on PBMC-EV interactions ex vivo

To this point, our *in vivo* and *ex vivo* results were obtained with Expi293F EVs that were separated using a standard workflow as previously published ^23^. To determine whether EVs from a different donor cell line, or obtained with a different separation protocol, would show a different distribution across blood compartments, we tested EVs from U-87 MG (glioma, “U-87”) cell culture that were separated by two methods: differential ultracentrifugation (UC) and size exclusion chromatography (SEC). Standard characterization was done for these EV preparations ^28^. Mean particle size and concentration and EV markers were comparable (Figure S5A, B, C), but cellular marker calnexin was more pronounced in UC-separated EVs (Figure S5B). Similarly, bovine serum albumin, a common co-isolate from growth media, was detected in UC-but not SEC-separated EVs. U-87 EVs were labeled with MemGlow and ultrafiltrated to remove unbound dye. We note that particle concentrations were lower after UF, as previously reported ^29,30^, and that the size distribution profile of MemGlow-U-87 EVs was narrower compared with unlabeled EVs (Figure S5D, E). U-87 EVs associated preferentially with CD20^+^ B cells at levels similar to those of Expi293F EVs (Figure 5A). Association of UC EVs was approximately 20% lower than for SEC EVs. Substantial variability for other cell types precludes firm conclusions except for the reproducible, much lower association of EV signal with non-B cell PBMCs (Figure 5B-F). Thus, EV from two donor cell types behaved similarly with regards to association with PBMC subtypes.

**Figure 5.**
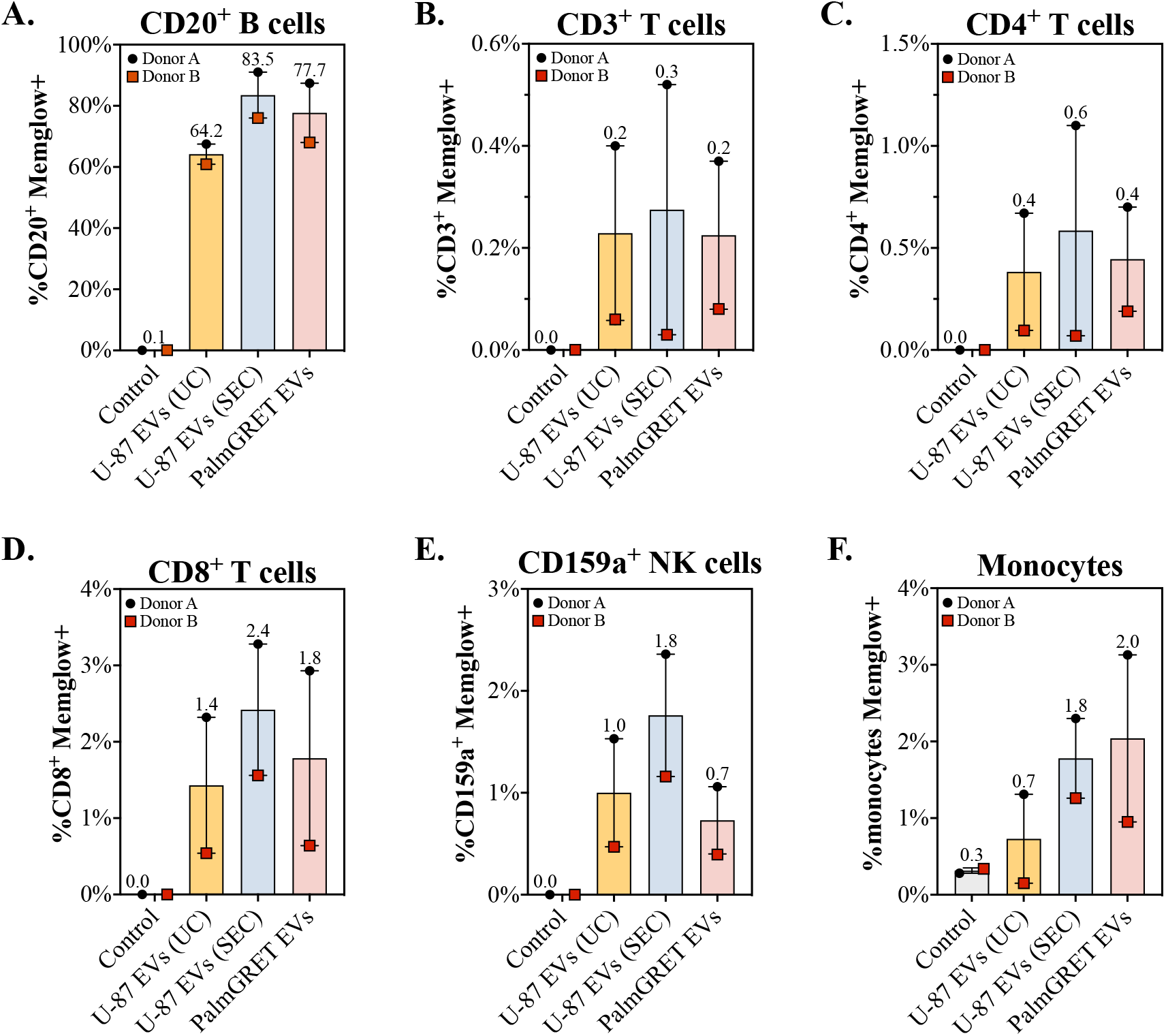
MemGlow-labeled EVs from U-87 MG cells preferentially interact with B cells *ex vivo*. Quantification of data obtained by flow cytometry using the same workflow described for Figure 3. Whole-blood was spiked with PBS MemGlow control, U-87 MG EVs labeled with MemGlow, or MemGlow-PalmGRET EVs from Expi293F culture and incubated for 5 min at 37°C. Percentage of MemGlow+ PBMC subtypes were determined with flow cytometry. The bars represent the mean of 2 independent experiments, performed with samples from 2 independent donors.

### Staining of PalmGRET-EV preparations with MemGlow alters EV GFP signal but not preferential associations between PalmGRET-EVs and B cells

Labeling of EVs with lipophilic dyes might change various properties of an EV preparation. The process of labeling with MemGlow appeared to decrease the percentage of PalmGRET EVs that were detected as positive for GFP (Figure S1C, E). Although this could be due to dye-mediated physical changes to the EVs, it might be explained most simply by formation of dye self-aggregates that decrease the EV share of the overall particle count. To understand if MemGlow labeling changed the cell association of EVs, 8E+08 PalmGRET EVs with or without MemGlow were tested in the *ex vivo* system. At 30 minutes, association was similar between the two populations of EVs and consistent with previous results (Figure S6).

### Influence of dosage on PalmGRET EV interactions with PBMC subtypes

Results of our previous time-course experiments suggest that EVs interact with PBMCs at levels detectable by flow cytometry as early as 5 minutes post-spike, peaking within 1 hour. We thus chose 30 minutes as the time point to interrogate the possible effect of EV dosage on cell associations, exposing blood from four donors to 8E+08 (dose 1), 4E+09 (dose 2), and 1.8E+10 (dose 3) PalmGRET EVs without MemGlow. There was a dose-dependent increase in GFP+ signal for all PBMC subtypes (Figure 6A). At the highest EV dose, all queried PBMC subtypes were on average >1% GFP+ (Figure 6 B-G). Importantly, monocytes disproportionately associated with EVs at the highest dose: on average, 26.2% of monocytes were GFP+ at dose #3 (Figure 6G). As a result, although CD20^+^ B cells were the predominant GFP+ cell type at dose #1 (as in previous experiments) (Figure 7A), the balance of GFP+ cells shifted to monocytes with increasing dose (Figure 7B).

**Figure 6.**
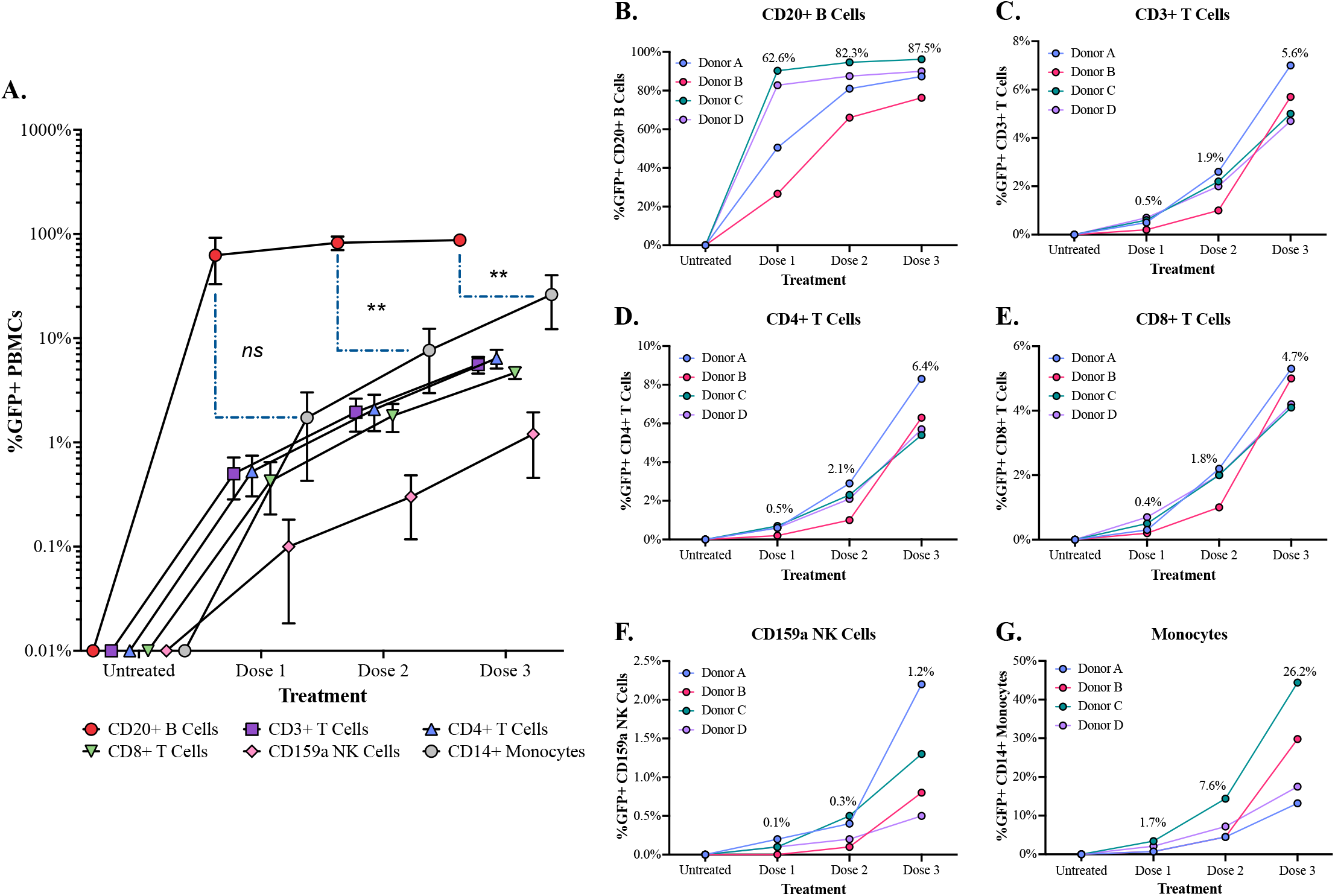
PalmGRET EVs preferentially interact with CD20^+^ B cells in an EV dose-response relationship 30 min after exposure to whole blood *ex vivo*. Cell-associated GFP signal was quantified by flow cytometry after 30 min of whole blood exposure to 3 doses of PalmGRET EVs: 8E+08 (Dose 1), 4E+09 (Dose 2), and 1.8E+10 (Dose 3). Data are expressed as % GFP+ PBMCs with all cell types and donor replicates plotted together (A) or separately (B-G) for (B) CD20^+^ B cells (C) CD3^+^ T cells, (D) CD3^+^CD4^+^ T cells, (E) CD3^+^CD8^+^ T cells, (F) CD159a^+^ NK cells, and (G). Curves in (A): arithmetic mean percentage of replicates (n = 4) and error bars: standard error of the mean. Connected symbols in (B-G) represent individual donors. Statistical comparisons for GFP+ CD20^+^ B cells and GFP+ monocytes at the same EV dose were performed using two-way ANOVA with Tukey’s post-hoc test, **p < 0.01.

**Figure 7.**
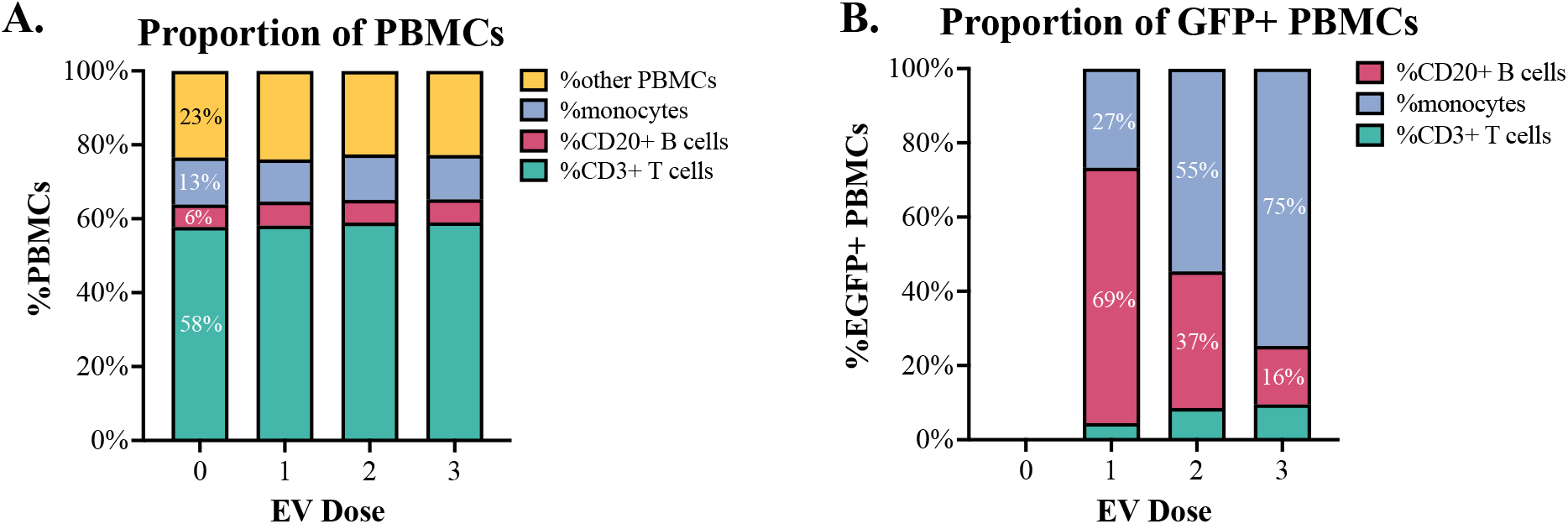
The dominant percentage of detectable EGFP+ PBMCs in PalmGRET-treated whole blood at different PalmGRET EV doses. (A) Comparison of PBMC subtype proportions and (B) proportion of GFP+ PBMC subsets (CD3^+^ T cells, CD20^+^ B cells, and monocytes). Graphs display mean frequency as a percentage of total PBMCs gated by PBMC subsets or GFP fluorescence. Data from experiments with blood of n = 4 donors.

## Discussion

Complementary to our recent *in vivo* biodistribution study, the present study demonstrated that *ex vivo* stimulation of whole blood with fluorescently-and genetically-labeled EVs is a reliable, sensitive, and physiologically relevant model suitable to study blood cell interactions with EVs. Previously, we demonstrated that EVs could be detected in association with CD20^+^ B cells within 1 minute of intravenous administration into the circulation of pig-tailed macaques. Remarkably, the *ex vivo* system used in this study also showed that EVs preferentially associate with CD20^+^ B cells in whole blood at levels similar to those detected *in vivo* in an EV dose-dependent manner. To the best of our knowledge, exposing whole blood *ex vivo* to labeled EVs as a means to quantify EV-PBMC interactions has not been performed previously.

Previous work by Fendl and colleagues established that the pre-analytical parameters chosen for collection and storage of whole blood can induce post-sampling release of EVs ^31^. Additionally, it has been well documented that different sample preparation methods can cause artifactual blood cell activation thereby altering cell phenotype, gene expression, and cell behavior^32–36^. Because experimental parameters likely influence how blood cells interact with EVs, the anticoagulant type, sample volume, handling, storage, transport, and incubation conditions should be carefully selected and compared. Blood cells are physically compacted in a confined space *in vivo* and changes to their environment can alter their cellular state and by extension interactions with neighboring cells and small particles, such as EVs. Shear force, blood flow, oxygen tension, three-dimensional architectural arrangement, and the presence of endothelial cells are parameters that are more technically challenging to recreate *ex vivo* but should also be considered, as they have been shown to influence the dynamics of blood cells in circulation *in vivo* ^37–39^. A potential strategy that could be explored in the future to examine the interaction network between EVs and blood cells involves the use of microfluidic devices. Due to recent advances in microfluidic technology which allow for tight control of system inputs, microfluidic devices can be customized to provide a continuous supply of oxygen, shear rate, and flow to blood samples to better represent physiological conditions ^40–45^. A comparison of these variables and techniques are warranted to inform future studies.

It was recently reported that labeling of PalmGRET-EVs with lipophilic dyes PKH26, DiD, and DiR caused an increase in EV size compared to their unlabeled EV counterparts ^46^. Changes in EV size are not inconsequential in biodistribution studies given that EV trafficking and uptake by recipient cells are partially determined by particle size ^47–49^. Here, we did not observe an increase in the mean size of MemGlow-PalmGRET EVs compared with PalmGRET-EVs. However, nano-flow cytometry analysis showed that MemGlow labeling of PalmGRET EVs decreased the % of GFP+ particles in PalmGRET EV preparations. It is unclear why MemGlow labeling would lead to a reduced percentage of GFP+ particles. The PalmGRET reporter anchors to the inner membrane leaflet of EVs ^11,46^, and the short, 12-carbon fatty acid tails of the MemGlow dyes ^50^ may label membranes without disturbing the inner membrane leaflet ^51^. Possibly, dye aggregates or dye interactions with non-EV substances may add to the total number of particles in the population. Nonetheless, no substantial differences were detected in terms of % GFP+ PBMC subsets when we compared the pattern of interactions between PBMCs and PalmGRET EVs labeled with or without MemGlow after co-incubation in whole blood *ex vivo* for 30 minutes at an EV dose of 8E+08. This suggests that both PalmGRET and MemGlow can be used reliably, either in combination or separately, to enumerate the interactions between blood cells and EVs *ex vivo*.

In the present study we detected a comparable degree of interactions between CD20^+^ B cells and EVs separated from the conditioned media of Expi293F-PalmGRET and U-87 cells. Expi293F-PalmGRET EVs and U-87 EVs isolated by SEC did not differ in terms of their preferential targeting of B cells. Moreover, U-87 EVs interacted at similar levels with CD20^+^ B cells, independent of the method of EV isolation. EVs produced by different cell types, including immune cells, stem cells, astrocytes, and cancer cells, often exhibit distinct biophysical properties and functions that typically align with those ascribed to their parental cells at the time of EV release ^52–57^. Nevertheless, EVs from two very different cell sources interacted with apparent preference with CD20^+^ B cells. It remains to be seen if this tropism is maintained by EVs from other cell types.

Our studies *in vivo* and *ex vivo* suggest that, at least for Expi293F and U-87 EVs, surface engineering of EVs may be dispensable for B-cell tropism in blood. Cellular tropism of EVs, including mechanisms of EV interactions with B cells, is an emerging area of interest ^58–60^. Previously, EVs have been surface-engineered with the major envelope glycoprotein of Epstein-Barr Virus (EBV), gp350, known to mediate attachment of EBV to B cells. On EVs, gp350 conferred B-cell tropism and cargo delivery functionality both *in vitro* ^61–64^ and *in vivo* ^64^. Platelet-derived EVs have also been shown to stimulate the production of antibodies from immortalized and primary B cells *in vitro*, although EV-B cell interactions were not specifically assayed ^65,66^. Microparticles derived from Kato cells interacted at high levels with B cells *in vitro* when the microparticles were preincubated with complement-containing human serum ^67^. Altogether, our studies and these previous findings support the notion that non-surface-engineered EVs may interact preferentially with B cells due to factors present in whole or fractionated blood. That is, preferential interactions with B cells occur primarily when EVs are isolated from blood samples, pre-incubated with blood serum, or added directly into whole blood samples *ex vivo*. Emerging evidence shows that when nano-sized particles, including EVs, are introduced into blood, they become coated with a “corona” ^68–71^ of proteins and other molecules that may influence particle-cell interactions. Possibly, non-surface-engineered EVs must adsorb plasma components to mediate interactions with B cells.

Although we found that B cells were the dominant PBMC subtype that interacted with EVs upon exposure to a low EV dose, interactions between EVs and monocytes at higher EV doses accounted for >50% of the cumulative GFP signal in CD3^+^ T cells, CD20^+^ B cells, and monocytes (Figure 7B). It is unclear if the dose-dependent shift in EV associations with PBMCs could simply be a result of enhanced recognition of EVs by phagocytic cells at high EV dosage or if it is due to a higher presence of phagocytic cells for clearance of EVs ^72–74^. Another possibility is that EVs saturate a specific receptor or receptors on CD20^+^ B cells in blood at a lower dose, leaving more unbound EVs to interact with other cell types such as monocytes. Learning more about the links between EV dosage and uptake, processing, and clearance mechanisms by phagocytic cells in blood will provide a basis to maximize EV circulation time and uptake by target cells for future clinical applications.

Since the flow cytometry methods we used here do not identify subcellular localization, our results are agnostic to the nature of EV-B cell association, i.e., surface interaction or internalization. We are currently addressing the subcellular fate of PBMC- and B cell-associated EVs in whole blood and whether these EVs influence cell functionality. However, both surface interaction and internalization are consistent with EV functions. Whereas internalization and membrane fusion are needed for cytoplasmic payload delivery, EVs may also communicate via surface-surface interactions that trip intracellular signaling pathways, thereby modulating recipient cell functionality.

*Ex vivo* models are useful to study EV interactions and functions in complex biological environments that are considered more physiologically relevant than simplified, monocultured single cell lines as *in vitro* models. Overall, this work supports the use of an *ex vivo* whole blood platform to interrogate PBMC-EV interactions in blood, which could help better understand the fate of intravenously injected EVs at the cellular and molecular level. While we limited this study to non-human primate whole blood, this *ex vivo* system could be easily tailored to study EV interactions with blood cells from other species. We fully expect this *ex vivo* model to serve as an intermediate step to investigate a broad range of physiological questions regarding the distribution and pharmacokinetics of labeled EVs in blood, while simultaneously eliminating the time, animal, and monetary costs of *in vivo* pharmacokinetics and biodistribution studies.

## Acknowledgments

The authors thank members of the Witwer Laboratory, the Retrovirus Laboratory, and Research Animal Resources for helpful discussions and suggestions.

## Supplementary Materials

**Figure S1.**
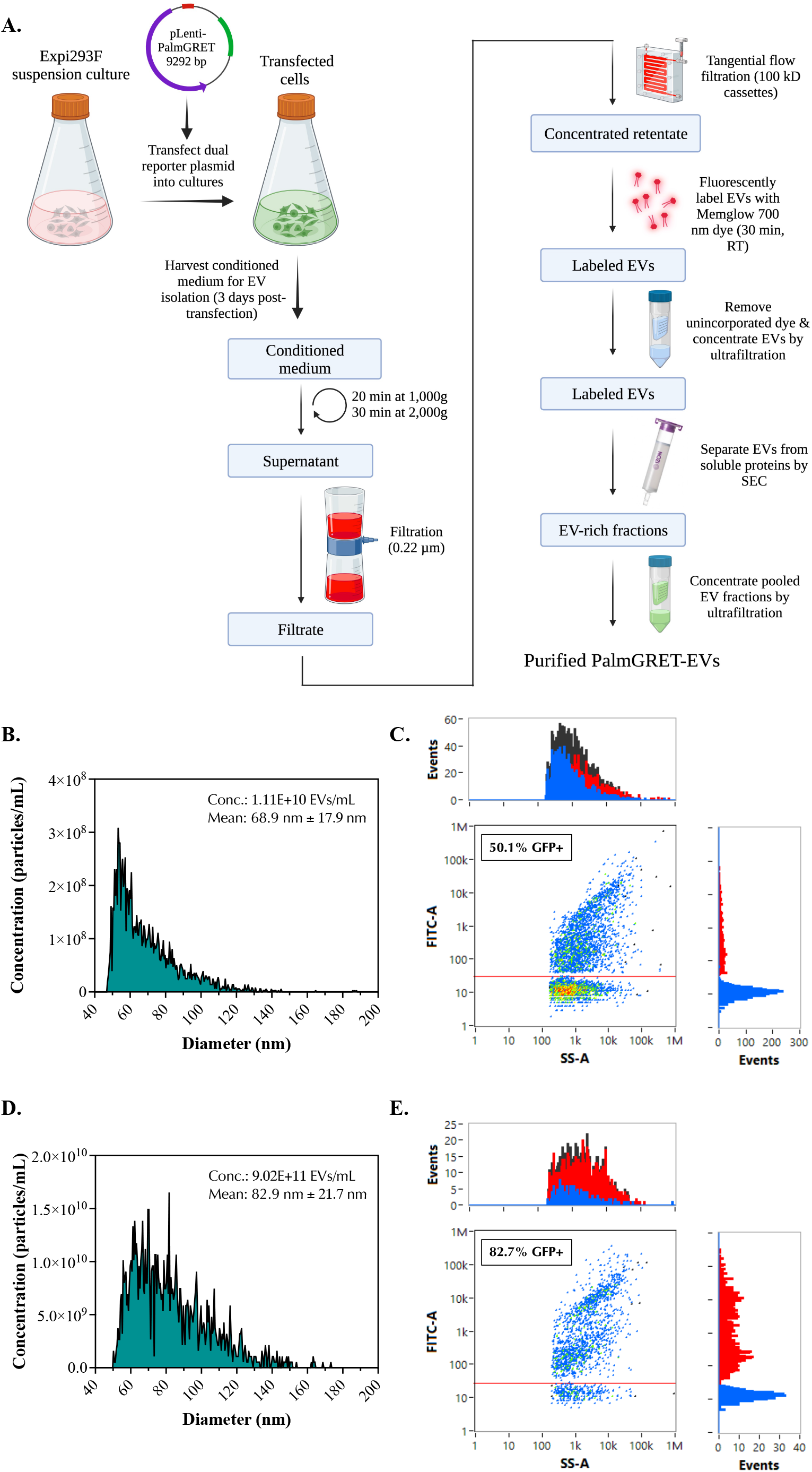
Isolation and characterization of EVs produced by PalmGRET-transfected Expi293F cells. (A) Schematic summarizing the workflow used to isolate EVs from large-scale cultures of Expi293F cells transfected with pLenti-PalmGRET dual reporter. Created with BioRender.com. (B-E) NFCM analysis of MemGlow-labeled and unlabeled PalmGRET EVs. (B,C) MemGlow-labeled PalmGRET EVs were diluted 50-fold in PBS and (D,E) unlabeled PalmGRET EVs were diluted 10,000-fold for analysis by NFCM. Histograms represent the particle size distribution and concentration of MemGlow-labeled (B) and unlabeled PalmGRET EVs (D). (C,E) Bivariate dot-lot plots of GFP fluorescence versus SS-A for EV samples analyzed by NFCM.

**Figure S2.**
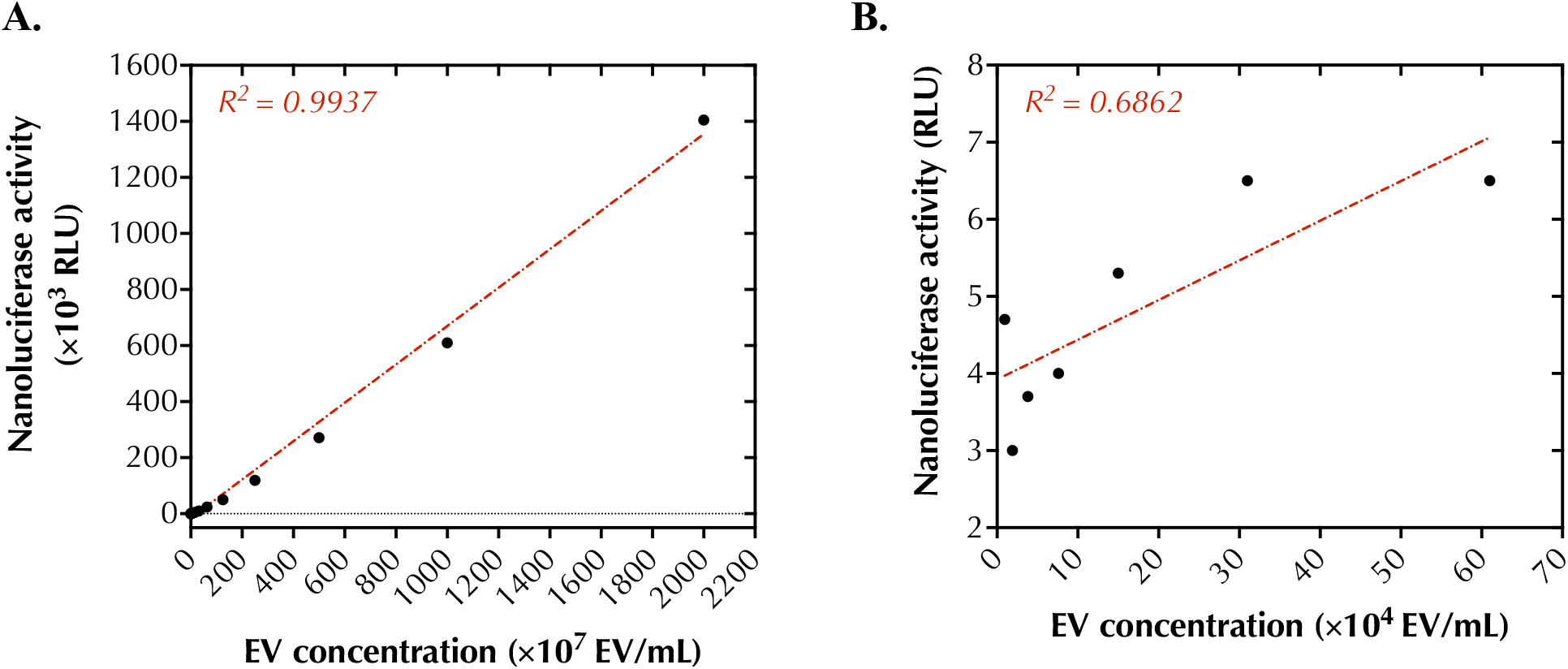
Relationship between particle concentration and nanoluciferase signal intensity. Two-fold serial dilutions of MemGlow-labeled PalmGRET-EVs in DPBS starting from a concentration of 2.0E+10 EV/ml were performed 22X to determine the limit of detection of PalmGRET-EVs in nanoluciferase assays. (A) A strong linear relationship between PalmGRET-EV concentration (EV/mL) and nanoluciferase signal (in terms of relative light units, RLU) was observed for the first 16 dilution samples (R^2^=0.9939). (B) EVs at a concentration lower than 1.64E+06 EV/mL exhibited a weaker correlation of particle concentration and nanoluciferase signal (R^2^=0.6842).

**Figure S3.**
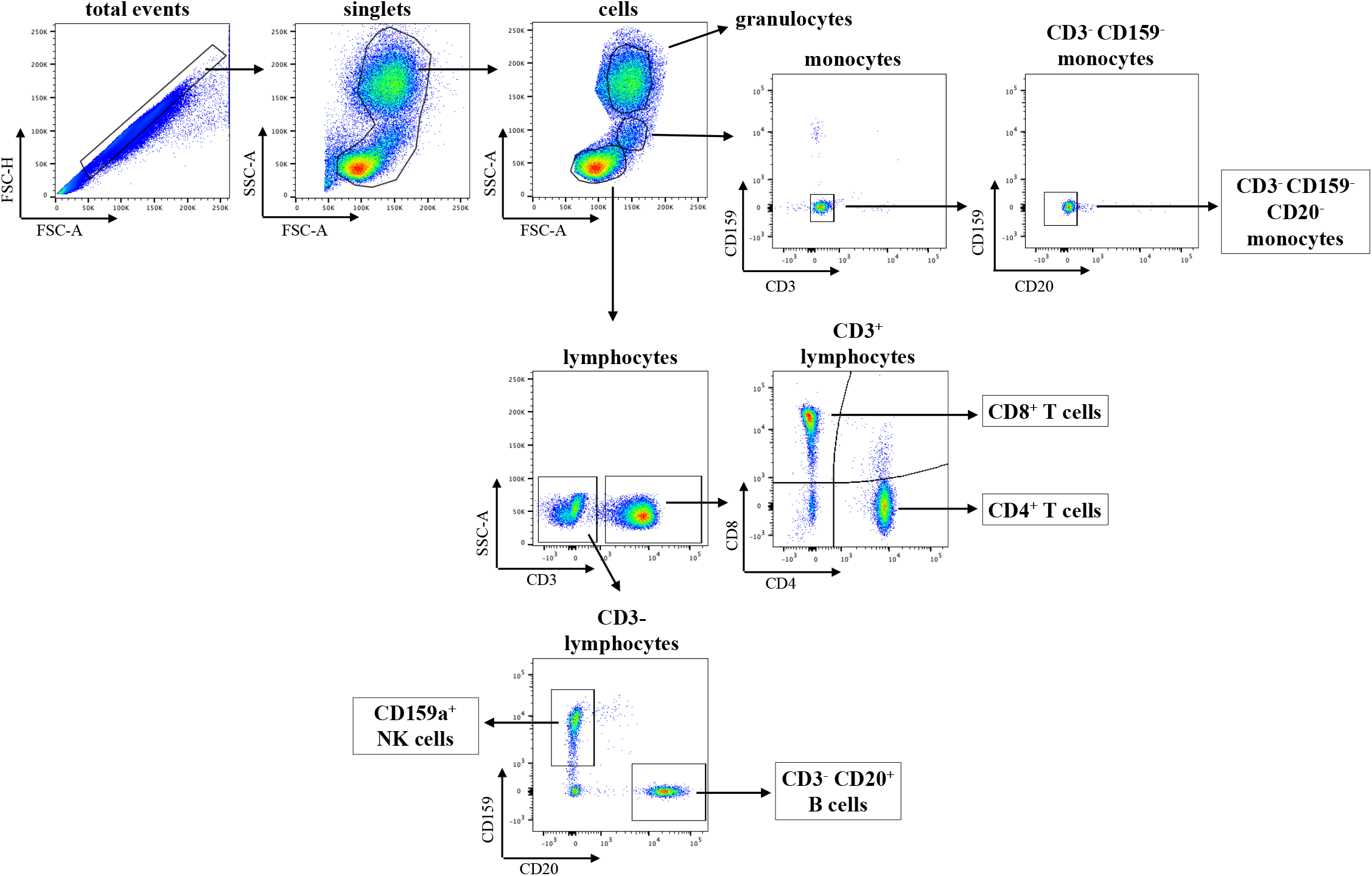
Representative flow cytometry plots of PBMCs showing the gating scheme to identify immune cell subtypes. Cells were identified by side scattering (SSC) and forward scatter (FSC) patterns. PBMC subtypes were identified using an antibody panel: monocytes (CD159-CD3-CD20- and CD14+ or CD14-), T cells (CD3+ and CD4+ or CD8+), B cells (CD3-CD20+), and NK cells (CD159+).

**Figure S4.**
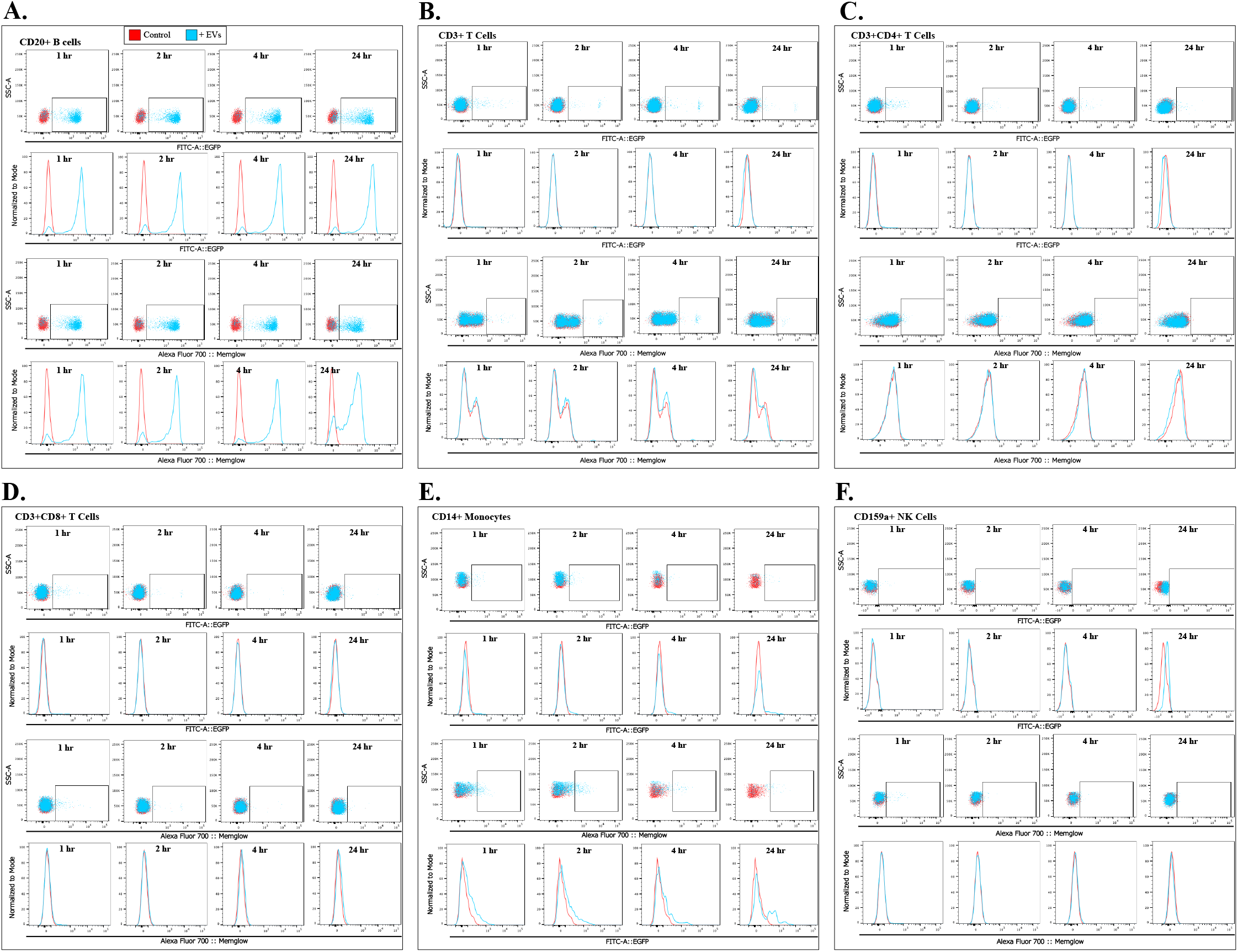
Representative flow cytometry plots of PalmGRET (EGFP)- and MemGlow-labeled EVs detected in association with PBMC subtypes after 24 h. FITC (GFP) and Alexa Fluor 700 (MemGlow) overlay flow cytometry dot plots and histogram plots of the major PBMC subpopulations from whole blood incubated with PBS (red data points & histograms) or MemGlow-PalmGRET EVs (blue data points & histogram) over 24 hours. EVs were detected based on GFP and MemGlow signal. Plots are representative of n = 2, whole-blood collected from 2 different pig-tailed macaques on the same day.

**Figure S5.**
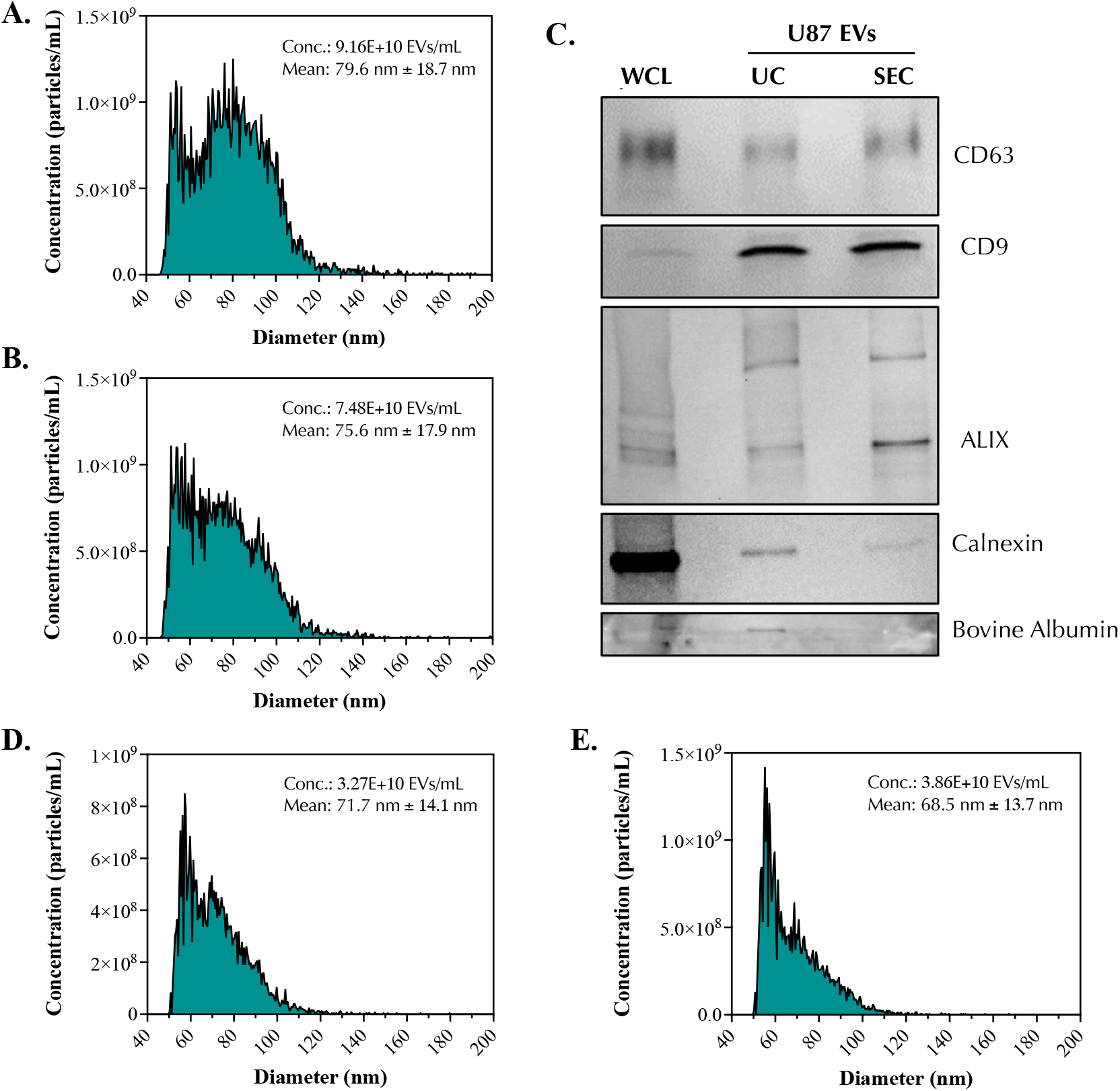
Characterization of U-87 MG-derived EVs. Nano flow cytometry measurement of particle size distribution and particle concentration of UC-purified U-87 EVs (A) and SEC-purified U-87 EVs (B). Western blot analysis of U-87 whole cell lysates (WCL), UC-purified U-87 EVs, and SEC-purified U-87 EVs (C). Size distribution and particle concentration profile of UC-purified U-87-EVs (D) and SEC-purified U-87-EVs (E) after MemGlow labeling.

**Figure S6.**
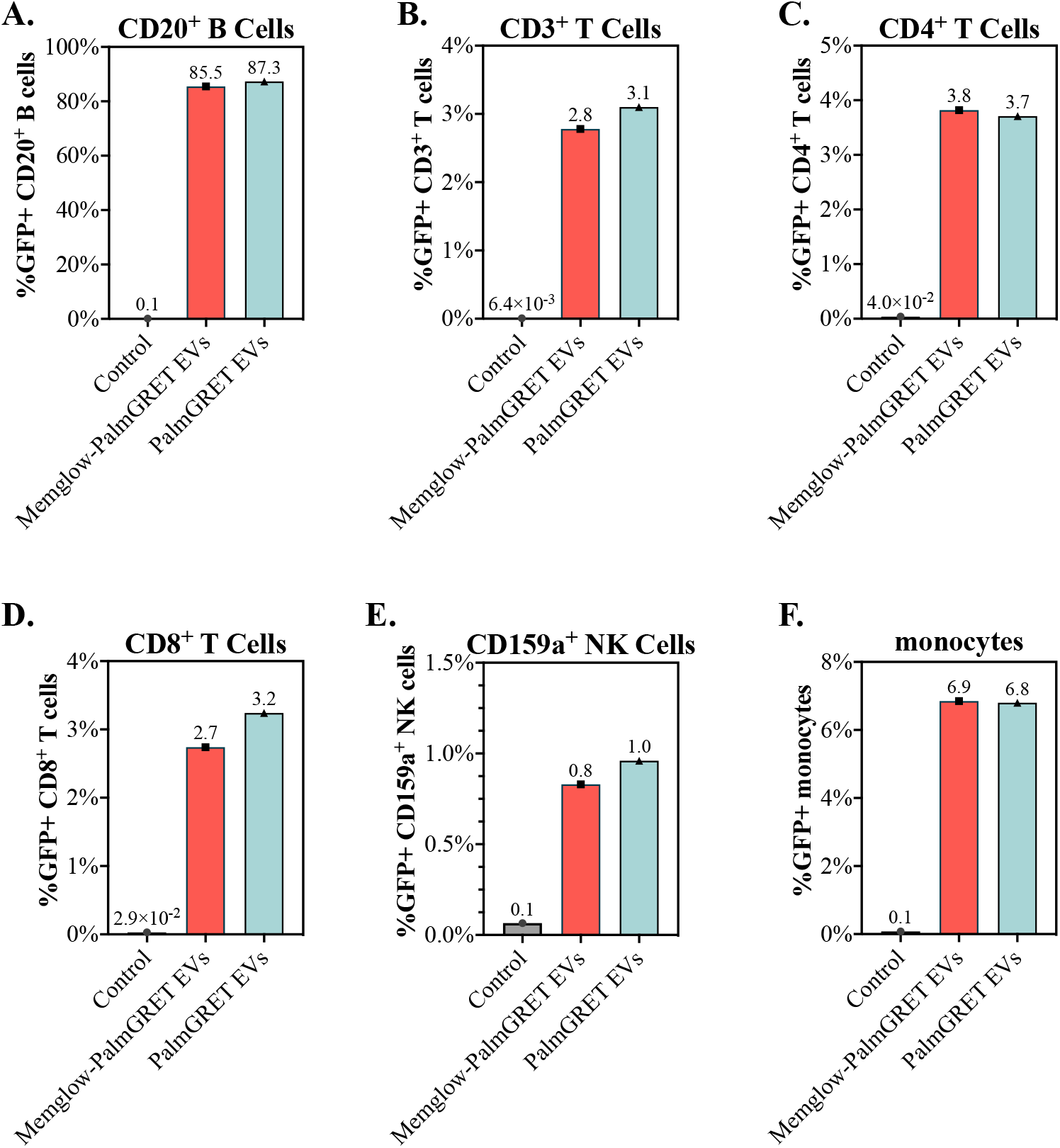
PalmGRET-EVs labeled with MemGlow interact with PBMCs similarly to unlabeled PalmGRET EVs. Flow cytometry Plots are representative of n = 1, whole-blood collected from 1 pig-tailed macaque.

